# Overexpression of Sirohydrochlorin Ferrochelatase Boosts Nitrogen Sulfur and Carbon Utilization in *Arabidopsis thaliana*

**DOI:** 10.1101/2024.03.22.586315

**Authors:** Naveen C. Joshi, Baishnab C. Tripathy

## Abstract

Nitrogen deficiency in the soil is a significant agronomic problem, and the application of nitrogenous fertilizers to the soil has environmental concerns, such as sodic soil and the emission of greenhouse gases. To increase the nutrient use efficiency, the cDNA of *AtSirB* coding for sirohydrochlorin ferrochelatase, responsible for Fe insertion to the tetrapyrrole moiety of sirohydrochlorin, was overexpressed in *Arabidopsis thaliana* under the control of 35S promoter for increased synthesis of siroheme. The siroheme is a cofactor for the plastidic enzymes nitrite reductase (NiR) and sulfite reductase (SiR), which reduces nitrite and sulfite to ammonium and sulfide, respectively. A three-step process including methylation, oxidation, and ferro-chelation produces siroheme from uroporphyrinogen III, an intermediate of chlorophyll (Chl) biosynthesis. The *NiR* and *SiR* gene expression and protein abundance increased in the over-expressers due to the increased AtSirB protein level. It resulted in an increase in N and S assimilation and enhanced protein content of over-expressers. Conversely, the total protein content decreased in antisense plants due to reduced NR and NiR activities. *AtSirB* over-expressers had higher protein and Chl contents and increased photosynthetic rate and biomass. Under N and S limitation, the protein, Chl, and photosynthetic electron transport rates in *AtSirB* over-expressers were higher than in WT. Results demonstrate that the *SirB* that hijacks uroporphyrinogen from the chlorophyll biosynthesis pathway is a crucial player in N and S assimilation. The siroheme is limiting for efficient nitrate and sulfate reduction and utilization. SirB could be genetically manipulated to increase crop productivity for sustainable agriculture.

## Introduction

Plants require an external source of nitrogen (N), which they obtain from the soil either as nitrate or ammonia. Application of nitrogenous fertilizers in the soil leads to the emission of the greenhouse gas nitrous oxide, which is responsible for global warming. Ammonia leached from soil causes eutrophication in water bodies (Anas et al., 2020; Stevens, 2019). Conversely, N deficiency negatively impacts crop growth and yield. Therefore, increasing N use efficiency will raise the fertilizer’s agronomic values by boosting crop yield, saving energy, and decreasing the negative environmental impact (Congreves et al., 2021).

After being absorbed from the soil, nitrate is transformed into nitrite by nitrate reductase (NR) using NADPH as a reductant (Campbell, 1999). Nitrite migrates from the cytosol to chloroplasts, where it undergoes six electron reduction mediated by plastidic nitrite reductase (NiR), often using ferredoxin as a reductant to form ammonium (Campbell, 1999). Similarly, a six-electron reduction of sulfite to sulfide is accomplished in the plastids of the photosynthetic organisms mediated by sulfite reductase (SiR) (Crane et al., 1996; Khan et al., 2010). Siroheme is the prosthetic group for both NiR and SiR (Murphy et al., 1974). It is a closed tetrapyrrole that contains Fe; its biosynthesis begins from uroporphyrinogen III, an intermediate of Chl and heme biosynthesis pathways (Murphy and Siegel, 1973). Three enzymatic reactions catalyze siroheme biosynthesis that includes (i) two steps of methylations, (ii) oxidation, and (iii) ferrochelatation (Wu et al., 1991; Tripathy et al., 2010). Precorrin-1 and precorrin-2 are produced due to methylation by Uroporphyrinogen III methyltransferase (UPM1) to synthesize siroheme (Warren et al., 1990). Precorrin-2 is dehydrogenated to form sirohydrochlorin. Sirohydrochlorin ferrochelatase (SirB) has a 2Fe-2S center and is the terminal enzyme in the siroheme biosynthesis pathway. It adds Fe to the center of sirohydrochlorin to produce siroheme (Raux-Deery et al., 2005).

Recently, research has focused on the effectiveness of nitrogen (N) consumption, including plant N absorption, acquisition, and utilization. Nitrate transporters are essential components of N homeostasis in plants. In the low-affinity nitrate transporter mutant *chl1*, overexpression of CHL1 led to the restoration of low and high-affinity nitrate uptake in *Arabidopsis* (Liu et al., 1999). In rice, over-expression of aspartate aminotransferase (AAT), a crucial enzyme in the production of amino acids, led to a noticeably greater level of leaf AAT activity and increased seed levels of both amino acids and proteins (Zhou et al., 2009). However, manipulating Nitrate reductase (NR) yielded a mixed result regarding N use efficiency, protein content, and photosynthetic efficiency. Constitutive expression of NR demonstrated a temporary delay in the loss of NR activity caused by drought, decreased nitrate content, and accumulation of high glutamine, which resulted in an increased amino acid pool in tobacco (Ferrario-me et al., 1998) and potato (Djennane et al., 2002). However, the total N content and protein content were unaltered. Similar efforts to overexpress the nitrite reductase gene in tobacco and *Arabidopsis* led to higher quantities of nitrite reductase transcripts but lower enzyme activity levels due to post-translational modifications (Crété et al., 1997). Although NiR enrichment enhances nitrite assimilation in *Arabidopsis*, NiR is posttranscriptionally controlled by nitrate or NR activity (Takahashi et al., 2001).

The glutamine synthetase (GS)/glutamate synthase (GOGAT) pathway is primarily responsible for ammonium assimilation in plants (Temple et al., 1998). Tobacco plants with overexpressed plastidic GS2 had higher photorespiration and better tolerance to high light intensity. Conversely, plants with GS2 were underexpressed, had low photorespiration, and were severely photoinhibited by high light (Kozaki and Takeba, 1996). Over-expression of chloroplastic GS2 in rice enhanced photorespiration and salt tolerance in transgenic plants (Hoshida et al., 2000). GS2 overexpression in tobacco was linked to both the drop in the leaf ammonium pool and the buildup of several free amino acids, like glutamate and glutamine (Migge et al., 2000). Reduced expression of Fd-GOGAT in a barley mutant showed a decrease in leaf protein and nitrate content (Häusler et al., 1994). In tobacco, overexpression of cytosolic pea GS1 improved plant growth and protein content (Fuentes et al., 2001). However, there are contradictory reports of the overexpression of GS1 and its impact on NUE. In tobacco (Chichkova et al., 2000) and poplar (Man et al. (2005), GS1 overexpression increased NUE, whereas overexpression of the soybean GS1 in pea plants did not increase the NUE (Fei et al., 2006). Even though all of these genes for secondary nitrogen assimilation seem to have the potential to increase NUE, the extent of their efficacy needs to be carefully investigated across various crops and cropping regimes. The effectiveness of genetically altering NR/NiR for significant increases in NUE met with only partial success and varies among plant species.

Sulfate assimilation by plants is necessary for life on Earth as it is an essential source for synthesizing the sulfur-containing amino acids cysteine and methionine (Tripathy et al., 2010; Garai and Tripathy, 2018). After being absorbed from the soil, sulfate is activated to become adenosine 5′-phosphosulfate (APS), and the primary assimilation process is the reduction of APS to sulfite (SO_3_^−2^), followed by its reduction to sulfide (S^−2^) (Saito, 2004). Plastid localized sulfite reductase (SiR) converts SO_3_^−2^ to S^−2^ via a six-electron reduction process mediated by the prosthetic group siroheme. Besides providing sulfide for the synthesis of cysteine, methionine, Fe-S centers, glutathione, and several other metabolic intermediates, SiR protects plants from sulfite toxicity. The expression of genes coding for NiR and SiR, responsible for N and S assimilation, is controlled by phytochrome (Faure et al., 1991; Sharma and Sopory, 1984; Sherameti et al., 2002). In addition, the expression of *SirB,* coding for sirohydrochlorine ferrochelatase (SirB) responsible for synthesizing the cofactor of siroheme that binds to both NiR and SiR, is also light-regulated. The promoter of *SirB* has light responsive elements (LREs) that photo-modulate *SirB* expression (Garai et al., 2016). Thus, the expression of both apoprotein and the cofactor of NiR and SiR is light-regulated. In *Arabidopsis thaliana*, the *AtSirB* [AT1G50170] codes for Sirohydrochlorine ferrochelatase, a nuclear-encoded enzyme with a 46 amino acid long transit peptide (Raux-Deery et al., 2005). The SirB is essential for plants as its knockout in *Arabidopsis* results in embryo lethality (Raux-Deery et al., 2005). The metabolisms of C, N, and S are interdependent in a significant way. N and S metabolism are often governed by nitrate and sulfate intake through the root system and their reduction to ammonium or sulfide.

Photosynthesis and respiration typically drive carbon metabolism. In this study, we have shown that the overexpression of AtSirB in *Arabidopsis thaliana* results in increased NiR and SiR expression, augmented NiR activity, higher protein content, and improved photosynthetic efficiency. In N and S limiting conditions, the over-expressers outgrow the wild type (WT) and have superior photosynthetic efficiency. Conversely, the *AtSirB* antisense plants either perished or had reduced nitrogen use efficiency, decreased protein content, lower photosynthetic efficiency, and were susceptible to N or S starvation.

## Results

### Generation and identification of *AtSirB* transgenic in *Arabidopsis thaliana*

A transgenic method was employed to characterize the function of Sirohydrochlorin ferrochelatase (AtSirB). Two constructs containing *AtSirB* cDNA in a sense (Overexpression) and antisense (Silencing) orientation with CaMV 35S promoter were transformed into *Arabidopsis* (Col-0) by *Agrobacterium*-mediated transformation **(Figure 1A)**. Over more than ten transgenic lines were generated. Using kanamycin-specific primers, PCR was used to detect transgenics. PCR findings showed that around 90% of the plants had incorporated the construct. Seeds were collected and stored individually from each transformed plant. Seeds harvested from each plant were selected in 30μg ml-1 kanamycin in 0.5X Murashige and Skoog medium (MS). All transgenic lines were segregated in a 3:1 ratio in a kanamycin plate, demonstrating a single integration of the transgene. Those who survived were grown for the next generation (T2). The resulting seeds were grown in kanamycin plates and were selected for kanamycin resistance. By employing 35S internal forward primer, gene-specific reverse primers (over-expresser), and gene-specific forward primers (antisense), gDNA PCR was used to verify T3 generations of homozygous transgenic plants **(Figure 1B).**

**Figure 1:**
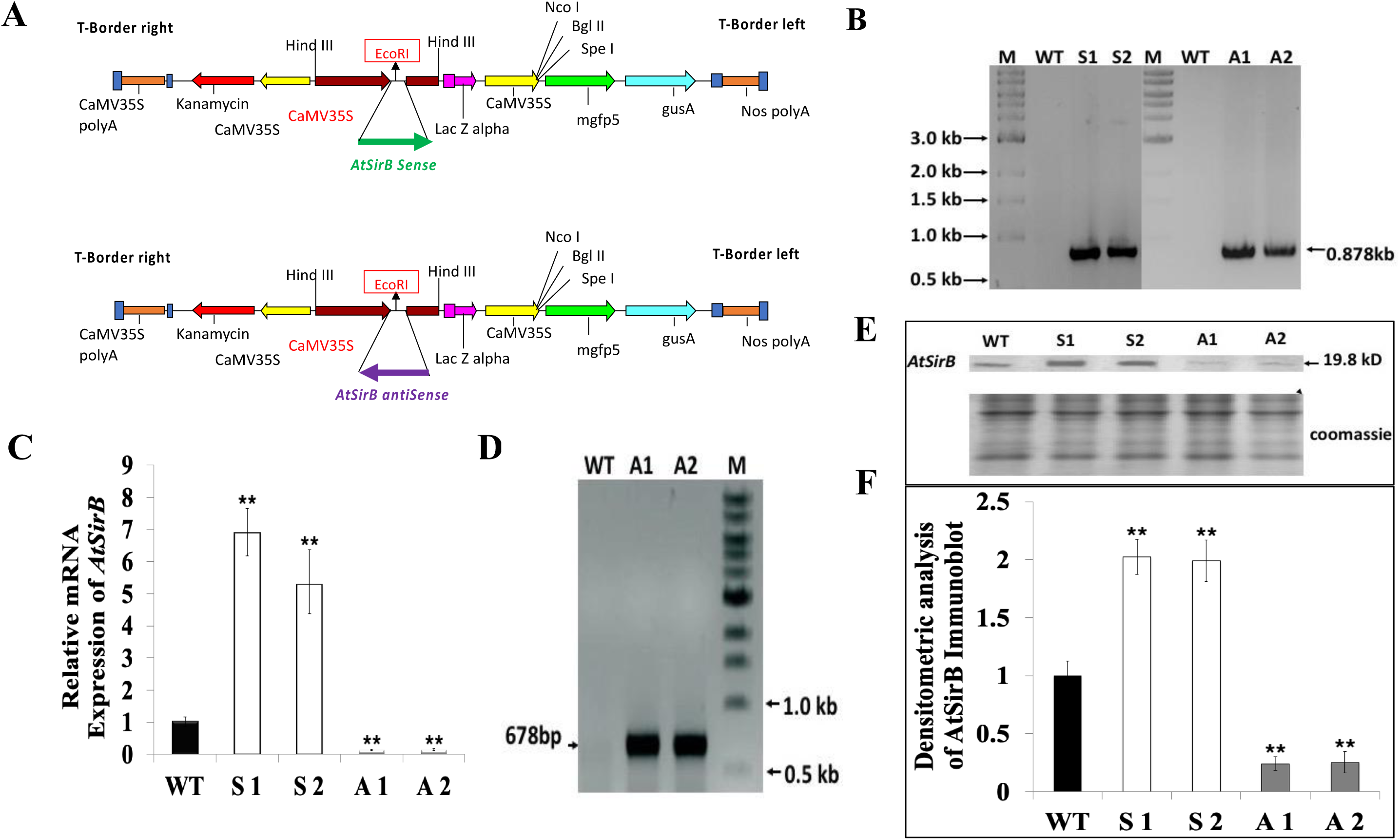
Transformation of Arabidopsis by using *AtSirB* cDNA. *AtSirB* cloned in pCAMBIA1304 modified plant transformation vector in sense (overexpression) and antisense orientation were used to transform *Arabidopsis* plants. (A) Vector constructs, (B) confirmation of T3 generation of *AtSirB* transgenic line by PCR by using gDNA as template and 35S forward primer and gene specific forward and reverse primers for antisense and sense plants, (C) Expression of *AtSirB* cDNA in transgenic over-expresser (S 1&S 2) and antisense (A 1&A 2) by quantitative real time PCR. (D) Antisense transgenic plants were further confirmed by antisense RNA analysis by strand specific RT-PCR, (E) AtSirB Protein expression in transgenic plants was checked by immunoblot against AtSirB antibody, Standard deviation is shown by the error bar. WT, S1, and A1 significantly differ from each other as indicated by the (*) asterisk (P< 0.05).

Once the transgene integration was confirmed by gDNA PCR, gene expression of *AtSirB* in WT and transgenic lines was checked by qRT-PCR analysis. RNA (2μg) extracted from WT and *AtSirB* transgenic plants were taken for cDNA preparation. Using the cDNA template, qRT-PCR was done by AtSirB gene-specific primers. The qRT-PCR reveals that, compared to WT, the AtSirB over-expresser lines (S1 and S2) had a 5-7-fold increase in *AtSirB* gene expression. In antisense lines (A1 and A2), *AtSirB* expression decreased by ∼70-80% **(Figure 1C)**. To probe further, the antisense *AtSirB* message abundance was studied. Since the forward primer only binds to antisense mRNA, it was possible to identify antisense mRNA using reverse transcription of RNA isolated from *AtSirB* antisense transgenic plants. The forward and reverse primers for *AtSirB* were used to create PCR targets from the first-strand reactions. The *AtSirB* was amplified by PCR using the cDNA prepared from the forward primer. The antisense A1 and A2 lines show a band corresponding to the AtSirB size of 678bp, which confirms the antisense expression. A cDNA prepared using total RNA from wild-type plants was used as a template that served as negative control **(Figure 1D)**.

Immunoblot analysis was conducted to study if the overexpression of AtSirB in transgenic plants resulted in an expected change in AtSirB protein expression. Total protein was extracted from 3-week-old WT, *AtSirBx* (S1, S2) and antisense (A1, A2) lines. The equal protein (20 mg) loading was confirmed by 12.5% SDS-PAGE. A polyclonal antibody preparation developed against *the Arabidopsis AtSirB* protein in our laboratory was utilized to immunodetect the AtSirB protein in *AtSirB* transgenic plants. Immunoblot showed that *AtSirB* protein expression was higher in S1 and S2 lines and lower in A1 and A2 transgenic lines. It proved that *AtSirB* was highly expressed in over-expresser plants and minimally in antisense plants **(Figure 1E-F)**.

### AtSirB modification changes plant morphology in *Arabidopsis*

The morphology of the *AtSirB* transgenic plants changed in a typical manner. The S1 and S2 plants were bigger and greener than the WT plants. In contrast, the antisense lines A1 and A2 survived and were smaller than WT **(Figure 2A).** Most of our antisense lines bleached and perished **(Figure 2B).** *AtSirB* transgenic root morphology changed, especially in S1 and S2 plants **(Supplementary Figure S 1A)**. Compared to WT, the number of lateral roots was higher in S1 and S2 plants. The number of lateral roots of antisense plants was the same as that of WT **(Figure 2C)**. In contrast to antisense plants, rosette diameter, fresh weight, and dry weight of S1 and S2 increased by 60%-80% **(Figure 2D, 2E, 2F)**.

**Figure 2:**
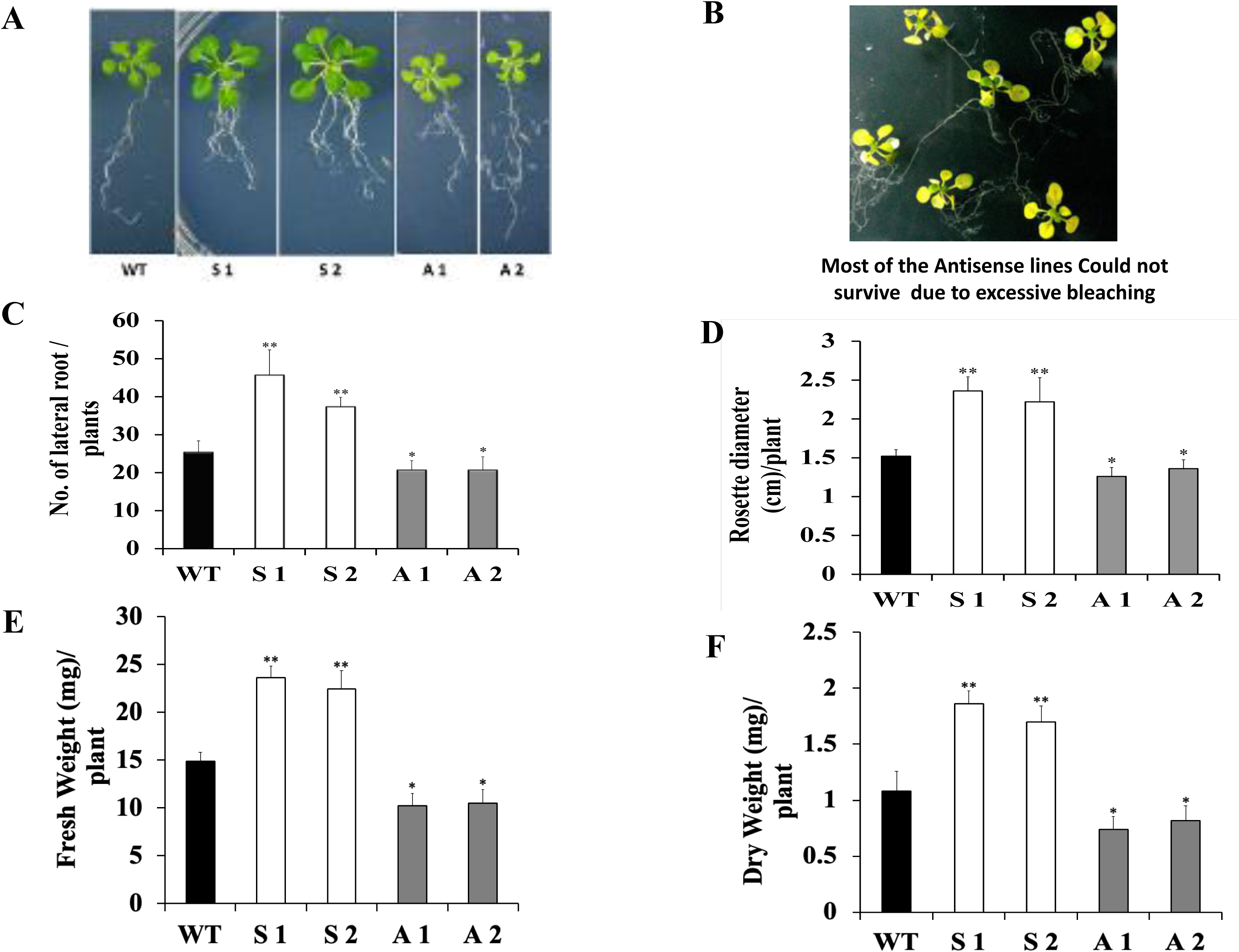
Phenotype and Morphological parameters *AtSirB* transgenic plants. Phenotype and morphological parameters of *Arabidopsis* transgenic lines. (A) Phenotypes of *AtSirB* sense (S 1 & S 2) and antisense (A 1 & A 2) lines grown for 3-weeks, (B) antisense lines which could not survive due to excessive bleaching, (C) lateral roots per plant, (D) rosette diameter, (E) fresh weight and (F) Dry weight. An average of 10 replicates makes up each data point. Standard deviation is shown by the error bar. WT, S1, and A1 significantly differ from each other as indicated by the (*) asterisk (P< 0.05).

### Modulation of N assimilation by *AtSirB*

SirB is the terminal enzyme responsible for the synthesis of siroheme, the cofactor of NiR. To probe if the increased expression of cofactor siroheme synthesizing gene *SirB* could modulate the expression of NiR apoprotein and NiR activity, its enzymatic reaction and expression were monitored.

### Gene expression and enzymatic activity of Nitrite Reductase (NiR) in *AtSirB* transgenic plants

The quantitative PCR of *NiR* (At2g15620) revealed that message abundance of *NiR* increased as a result of *AtSirB* overexpression. The qRT PCR indicated that the *NiR* gene expression in the overexpression lines, S1 and S2, was 90 % and 80 % higher than WT. However, in antisense lines (A1 & A2), there were no significant changes in the gene expression of NiR **(Figure 3A)**. Due to increased message abundance for NiR, the NiR apoprotein increased in S1 and S2 over-expresser lines. The Western blot further revealed that in A1 and A2 antisense lines, there was no significant change in NiR protein level **(Figure 3B)**. The NiR enzymatic activity was measured to examine the consequences of the altered expression of *SirB* in transgenic plants. Four-week-old WT and *AtSirBx* and antisense plants, grown at 21^0^C at 100μmol photon m-2s-1 and 14h L/10h D photoperiod, were taken to estimate NiR activity. Compared to WT, NiR activity increased by approximately 20%-25% in S1 and S2 lines due to the increased availability of the cofactor siroheme. However, in antisense plants, its enzymatic activity declined by 50%-60% **(Figure 3C)**. In S1 and S2 overexpresser plants, higher activity of NiR efficiently consumes its substrate NO_2_^−^ whose over-accumulation could be toxic to the plant.

**Figure 3:**
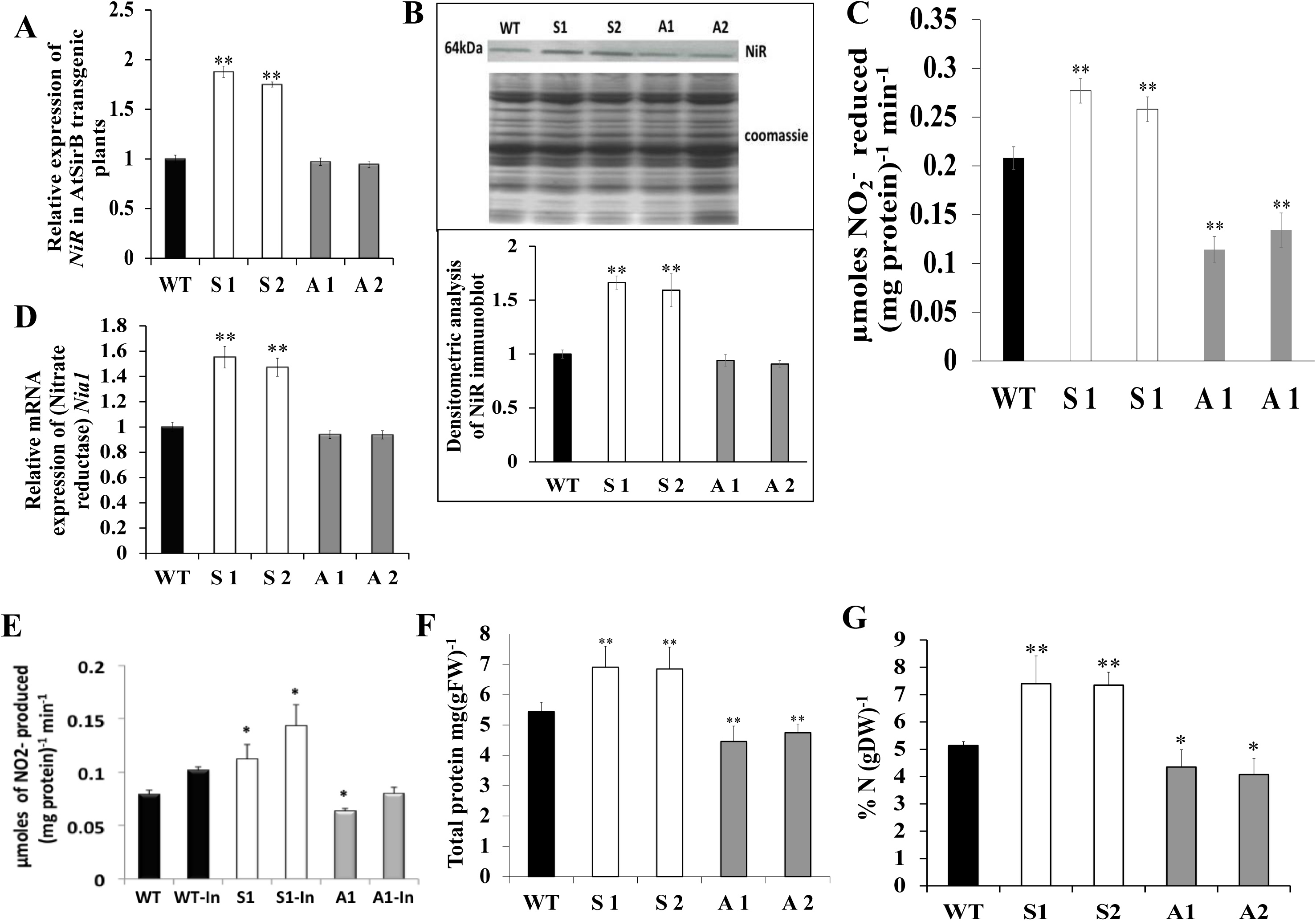
Characterization of *AtSirB* transgenic plants. WT and *AtSirB* sense lines (S1, S2) and antisense lines (A1, A2) plants raised at 22^0^C under 14h L / 10h D photoperiod in a light intensity of 100 µmoles photons m^−2^ s^−1^ for 3 weeks in MS medium and their vital parameters were analysed. (A) Quantitative PCR of gene expression of nitrite reducatse (NiR), (B) NiR protein abundance using immunoblot, (C) nitrite reductase (NiR) activity estimated in the leaves of 3-week-old WT and transgenic plants, (D) Nitrate reductase *(Nia1)* gene expression was quantified by qRT-PCR, (E) nitrate reductase activity was measured in leaf of 3-week-old WT and transgenic plants; when required, plants were initially treated with 3mM of KNO_3_ to induce (In) NR, (F) total protein content of *Arabidopsis* seedling, (G) nitrogen content was quantified by CHNS analyzer using L-Cysteine as standard. An average of 5 replicates make up each data point. The standard deviation is shown as the error bar. WT, S1, S2, A1 and A2 significantly differ from each other as indicated by the (*) asterisk (P< 0.05).

### Gene expression and enzymatic activity of Nitrate Reductase (NR) in transgenics

The cytoplasmic NR, which converts NO_3_^−^ to NO_2_^−^, supplies the substrate for NiR. Consequently, their stoichiometry needs to be regulated. To probe further, transcript abundance and NR activity in WT and transgenic lines were examined. In S1 and S2 over-expresser lines, the gene *NIA2*, which codes for NR, showed increases in expression of 60% and –50%, respectively **(Figure 3D).** To ascertain if NiR could modulate NR, its activity was estimated in 3-4-week-grown WT and *AtSirB* transgenic plants (S1 and A1). NR activity in S1 plants increased by ∼39 % over WT. Conversely, A1 plants’ activity declined by ∼20 % **(Figure 3E)**.

### *AtSirB* overexpression increased total protein and N levels

#### Total protein content

The *AtSirBx* plants had higher levels of expression and activity of enzymes involved in N assimilation. As a result, the total protein content of transgenics increased. Compared to WT, S1 and S2 had a 30% and 28% higher protein content, respectively. In antisense plants, enzymatic activities of NiR and NR were substantially reduced, likely due to siroheme limitation. It resulted in lower protein content (15%-20%) of antisense plants **(Figure 3F)**.

#### N content

The significant increase in the enzymatic activity of two crucial enzymes involved in N metabolism and total protein contents led us to quantify total N content in WT and *AtSirB* transgenic plants. N content increased by ∼31% and 30% in S1and S2 overexpressor and declined by ∼16% and 15% in A1and A2 antisense plants compared to WT **(Figure 3G)**. It is in agreement with that of the total protein contents.

### *AtSirB* manipulation modulates photosynthetic efficiency in *Arabidopsis*

#### *AtSirB* manipulation alters pigment Contents in *Arabidopsis*

WT and *AtSirB* transgenic plants (S1, S2 and A1, A2) were grown in MS plate for three weeks. Their total Chl contents were measured. S1 and S2 had higher amounts of total Chl (∼31%) than WT. Compared to WT plants, overall Chl content decreased (∼30%) in A1 and A2 plants **(Figure 4A)**.

**Figure 4:**
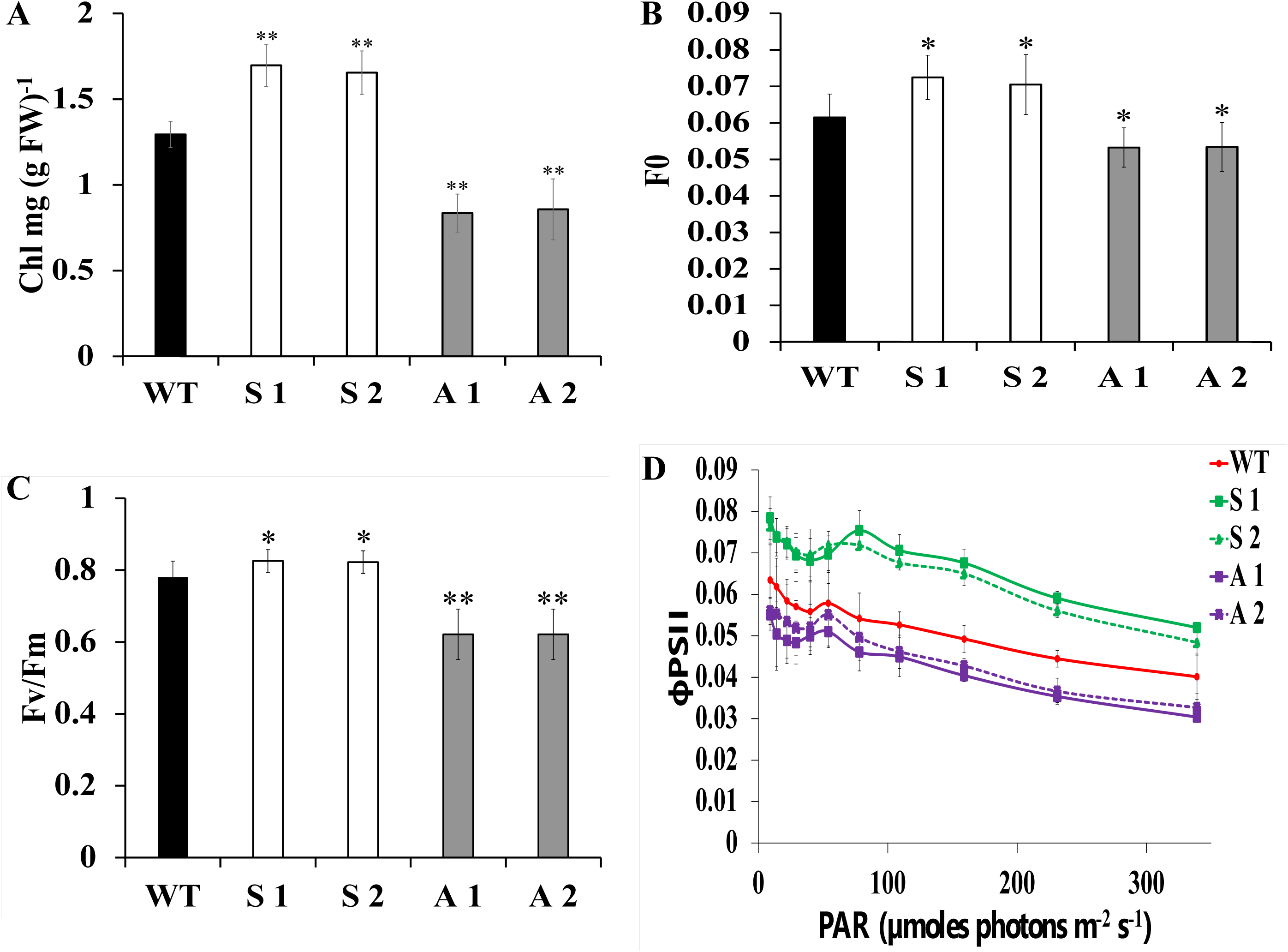
Photosynthetic efficiency of WT and *AtSirB* transgenic plants. WT and *AtSirB* transgenic sense (S1) and antisense (A1) plants were grown at 22^0^C under 14h L / 10h D photoperiod in a light intensity of 100 µmoles photons m^−2^ s^−1^ for three weeks in MS medium and their (A) Chl, (B) F0, (C) Fv/Fm, and the (D) фPSII were measured. An average of 10 replicates makes up each data point. Standard deviation is shown as the error bar. WT, S1, and A1 significantly differ from each other as indicated by the (*) asterisk (P< 0.05).

### *AtSirB* transgenic accumulates Chl biosynthetic intermediates in *Arabidopsis*

To study the Chl biosynthetic potential of WT, S1 and A1, the contents of Pchlide and Proto IX, the intermediate of Chl biosynthesis pathway, were estimated by spectrofluorometry (Hukmani and Tripathy, 1993). Under steady-state illumination (100 μmol photon m-2s-1), compared to WT, the amounts of Pchlide were augmented by 53% in S1 and declined by 37% in A1 plants **(Supplementary Figure S2A)**. Like Pchlide, the protoporphyrin IX content of S1 plants increased by 34%, whereas in A1 plants, it decreased by 27% **(Supplementary Figure S2B)**.

### Changes in Chl a Fluorescence parameters due to the genetic manipulation of *AtSirB*

Chl fluorescence is a non-invasive marker of photosynthetic ability in plants (Padhi et al., 2021). In order to ascertain whether plants with higher Chl and protein concentrations had higher photosynthetic ability, three Chl a fluorescence metrics, Fo, Fv/Fm, and the quantum yield of PSII (фPSII), were measured. The Fo, the minimal fluorescence increased in S1 and S2 (sense lines), likely due to an increase in Chl content. In the antisense lines (A1 and A2), Fo declined, likely due to a reduction in Chl content **(Figure 4B).** The maximum photochemical efficiency of the dark-adapted leaves, measured as Fv/Fm ratio, significantly increased in S1 plants compared to WT plants. In contrast, it decreased by almost 20% in A1 plants **(Figure 4C)**. As expected, the фPSII, the quantum yield of PSII in light-adapted plants, reduced in response to rising light intensities in WT and transgenic plants. In saturating light intensities, the фPSII of S1 and S2 lines were 40 % and 32 % higher than WT, respectively. Conversely, in the antisense A1 lines, the фPSII was substantially lower than WT at saturating light intensities **(Figure 4D)**.

### *AtSirB* expression in N-limitation condition

To ascertain if *SirB* overexpression could mitigate N deficiency, the NO_3_^−^ content in the growth media was reduced, and the WT and transgenic plants were cultivated under N-limiting conditions. WT and transgenic plants were first grown for 15 days in a regular MS medium before moving to N-limiting media in agar plates with Hoagland medium adjusted to 0.1N NO_3_^−^ concentrations. After seven days in the N-limiting medium, there was a visible difference in *AtSirB* transgenic plants compared to WT. The phenotype of WT plants under 0.1N condition appears pale due to N deprivation. The S1 plants appeared greener than WT when grown in identical growth conditions. However, in A1 plants, the N starvation caused severe phenotypic changes, including bleaching **(Figure 5)**. Total Chl contents were higher in S1 plants than WT and lower in A1 plants at all N levels. The Chl contents of S1 were 87% higher than WT at 0.1N levels. The Chl content decreased by 30% compared to WT at 0.1N level in A1 plants **(Figure 6A)**. Total protein content was higher in S1 and lower in A1 plants when compared to WT. Under normal (1N) N level, protein contents of S1 were 35% higher, and in A1, the same was lower by 11% than that of WT. The protein contents of S1 plants were 95% higher than WT’s in 0.1N levels. In A1 plants, protein contents declined by 32% compared to WT **(Figure 6B)**. To comprehend the mechanism underlying the tolerance of *AtSirB* transgenic plants to N deprivation, the activities of N assimilating enzymes were investigated in WT and transgenic plants grown in standard and N-limiting medium, compared to WT, NiR activity at optimum N level (1N) increased by 30% in S1 plants. In A1 plants, the same declined by 45%. The degree of NiR activity augmentation in S1 rose in N-limiting conditions compared to optimal N levels, whereas it reduced in A1 plants. S1 plants had NiR activities that were 72% greater in 0.1N levels than WT plants. In the A1 plants, NiR activity dropped by 54% compared to WT **(Figure 6C)**.

**Figure 5:**
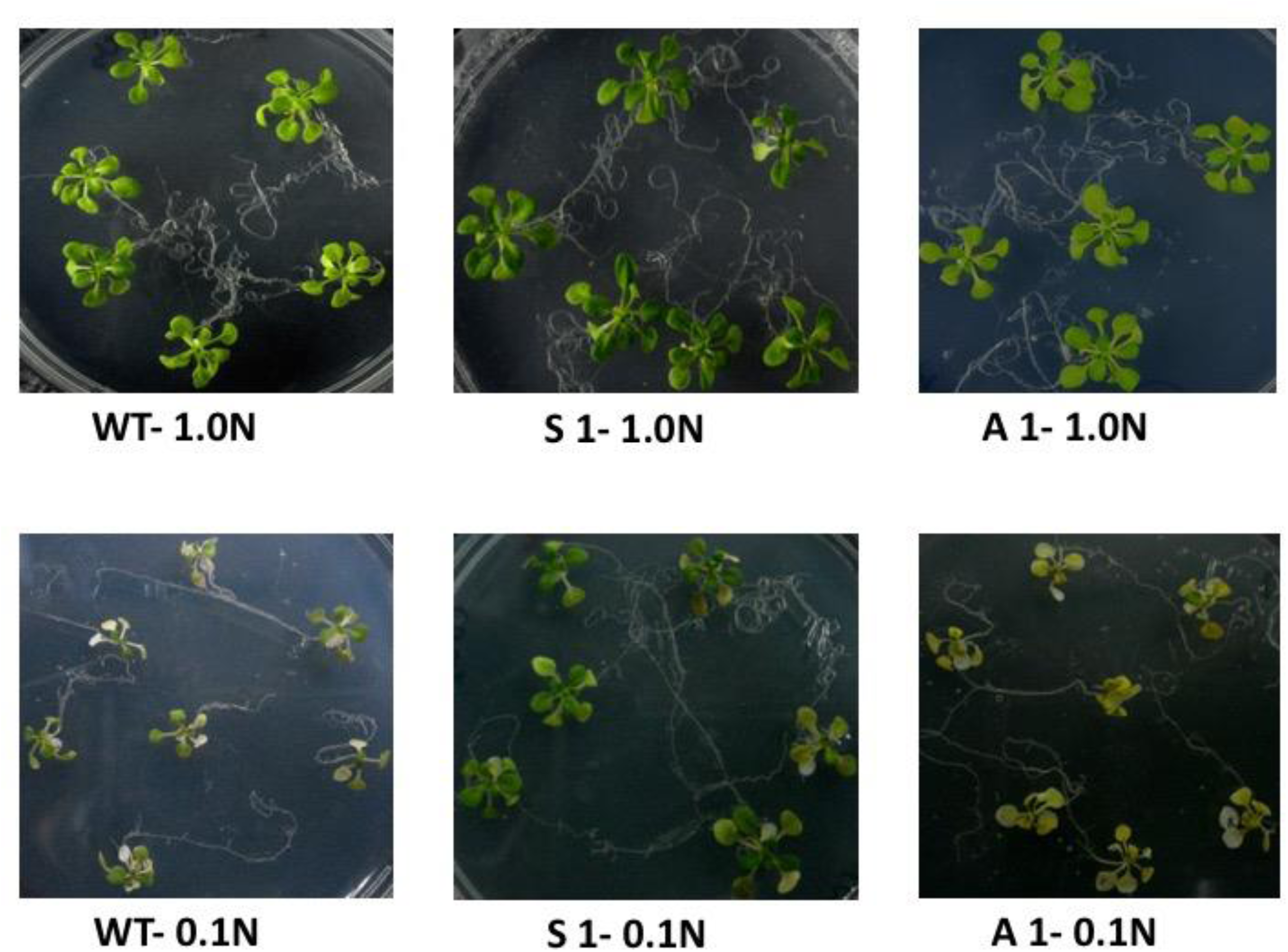
Phenotypes of WT and *AtSirB* transgenic plants in N-limiting condition. Plants were grown at 22^0^C under 14h L / 10h D photoperiod in (100 µmoles photons m^−2^ s^−1^). WT and *AtSirB* transgenic plants (S 1 and A 1) raised under normal (1.0N) MS for 15 days and subsequently transferred for 7 days to Nitrogen limitation medium (0.1N). Nitrogen concentration for 1.0 N was 5mM.

**Figure 6:**
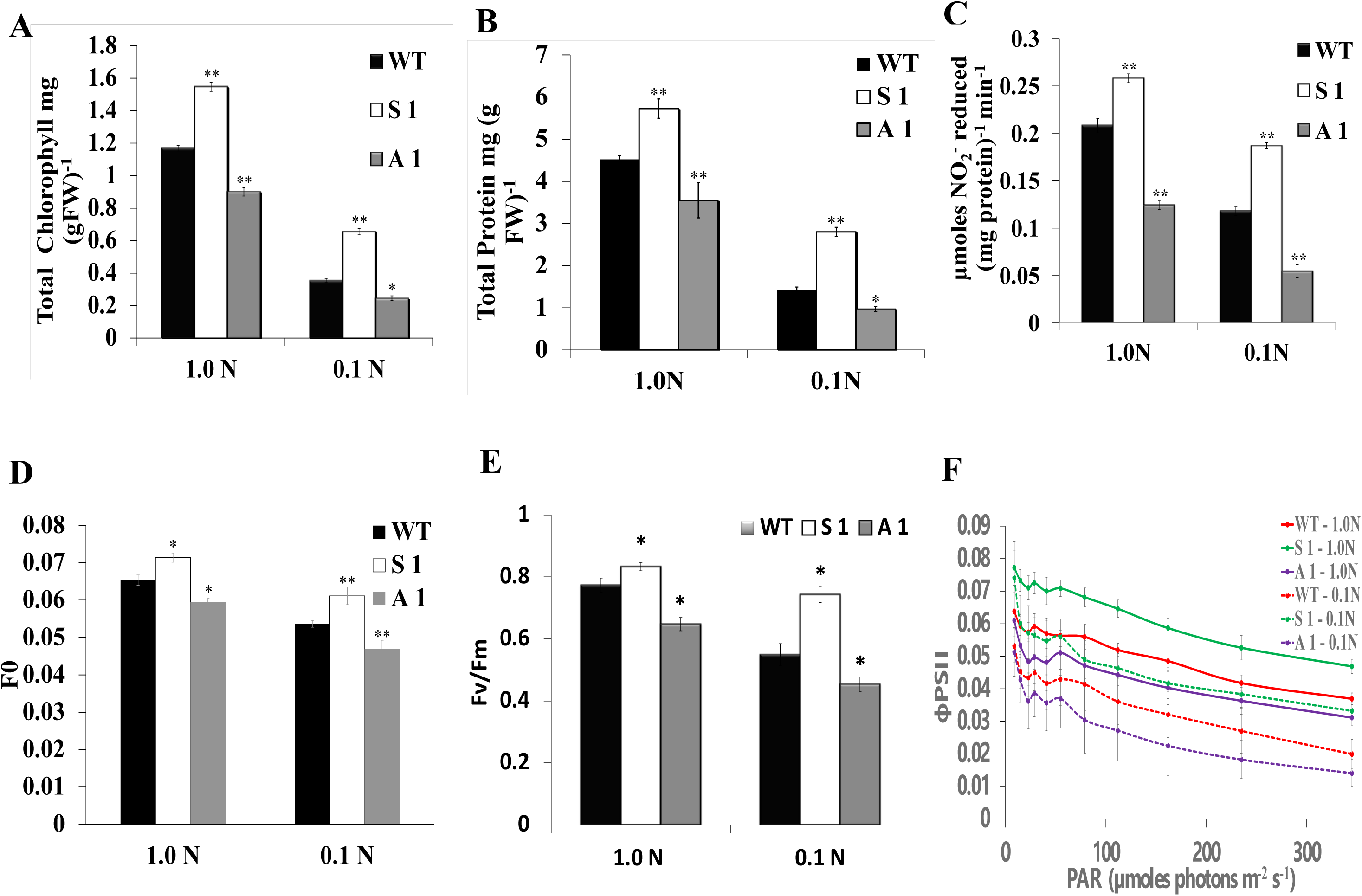
Responses of WT and transgenic plants to N-limitation. WT and *AtSirB* transgenic plants (S1 and A 1) were grown at 22^0^C under 14h L/10h D photoperiod in a light intensity of 100 µmoles photons m^−2^ s^−1^ in MS for 15 days and subsequently sifted for 7 days into nitrogen limitation condition (0.1N) and their physiological parameters were monitored. (A) Total Chl, (B) total protein, (C) NiR activity, and (D) Fo, (E) Fv/Fm, (F), фPSII measured by pulse amplitude modulated (PAM) fluorometer. An average of 10 replicates makes up each data point for A-C and 10 replicates for D-F. The standard deviation is shown by the error bar. WT, S1, and A1 significantly differ from each other as indicated by the (*) asterisk (P< 0.05).

### Chl a fluorescence parameter in WT and transgenics grown in optimal and N-limiting conditions

WT and transgenic lines were grown for 15 days in regular MS medium before being shifted to agar plates with Hoagland medium. The N level of the Hoagland medium was adjusted to 0.1N of the normal concentrations by reducing NO_3_^−^ concentrations. F0, Fv/Fm, and фPSII were examined after seven days of growth on N-deficient media. F0 declined in nitrogen-limiting conditions in WT, S1, and A1 plants, although S1 plants had high F0 in comparison to WT, whereas A1 had lower F0 than WT at all nitrogen levels **(Figure 6D)**. In comparison to WT plants grown at all N levels, S1 plants had a higher Fv/Fm ratio. Conversely, in A1 plants, the same declined **(Figure 6E)**. Both in N-sufficient and N-limiting conditions, the фPSII increased in response to increasing light intensities. In *SirBx* plants, the фPSII was higher in S1 in limiting and saturating light intensities. The higher light-saturated rate implies an increase in the antenna size and the components of the electron transport chain in *AtSirBx* plants, and the increased slope of фPSII at limiting light intensities supports enhanced energy collection by the light-harvesting complex. In N-deficient media, the фPSII was higher in overexpressers and lower in antisense plants **(Figure 6F)**.

### Alteration of carbohydrate metabolism in WT and *AtSirB* transgenic plants

Sugars and starch build up in leaves due to an N deficit as the carbon skeletons cannot be utilized for amino acid and protein synthesis. WT and *AtSirB* plants were cultivated in regular MS medium for 15 days before being sifted to agar plates with Hoagland medium lowered to 0.1N of the usual concentrations by lowering NO3-concentrations for seven days. WT and *AtSirB* transgenic plants were stained with iodine to show the variations in starch distribution following seven days of growth in N shortage conditions. No starch accumulation was observed in WT, S1, and A1 plants in optimal N concentration. Under N-limitation (0.1N), carbohydrates accumulated in the leaves of WT and A1 plants. However, the S1 plants did not accumulate carbohydrates when grown in a 0.1N growth medium, suggesting that they could tolerate extreme N-starvation **(Supplementary Figure S3)**.

### Modifications of *AtSirB* expression change sensitivity towards S limitation condition

To investigate the role of *AtSirB* in S metabolism, WT and *AtSirB* plants were grown in S-limiting media by reducing the SO_4_^−2^ level of the growth medium. WT and *AtSirB* plants were first grown for 15 days in regular MS medium before being transplanted to agar plates with Hoagland media with a normal or limiting S concentration (0.1S). Compared to WT plants, *AtSirB* transgenic plants showed a noticeable change in phenotype after seven days of growth on S-deficient media. The phenotype of WT plants looked pale-green after seven days of growth in S starvation (0.1S) media. The S1 plants seemed greener than WT under the same growth conditions. However, the S deprivation of A1 plants resulted in significant phenotypic alterations, and the plants became severely blanched **(Figure 7A)**. As the siroheme is a prosthetic group of *SiR* (At5g04590), its expression was studied in optimally grown (1S) WT, *AtSirBx* (S1, S2) and *AtSirB* antisense (A1, A2) plants. Compared to WT*, sulfite reductase (SiR)* expression increased by 55% and 50% in S1 and S2 plants, respectively. In A1 and A2 plants, SiR message abundance was slightly lower than the WT **(Figure 7 B)**. The protein abundance of SiR was examined by Western blot to see whether gene expression and protein expression are correlated. In consonance with gene expression, the S1 and S2 lines displayed a 50% and 60% increase in SiR protein abundance over the WT. However, their abundance in A1 and A2 lines was similar to WT **(Figure 7C)**. To probe further, WT and *AtSirB* transgenic plants grown in optimal (1S) and minimal (0.1S) S growth media and their Chl contents were measured. In S1 plants grown in optimal S medium (1S), total Chl levels were higher (32%) than those of WT plants. However, in A1 plants, Chl content declined by 27% **(Figure 8A)**. Similarly, the S1 plants had a 28% higher total protein content than WT plants in optimal growth conditions (1S). Conversely, protein content was reduced (29%) in antisense A1 plants. In response to S-starvation (0.1S), the protein content of S1) was 46% higher than that of WT. The protein content declined by 51% in A1 plants grown in an S-deficient medium **(Figure 8B)**.

**Figure 7:**
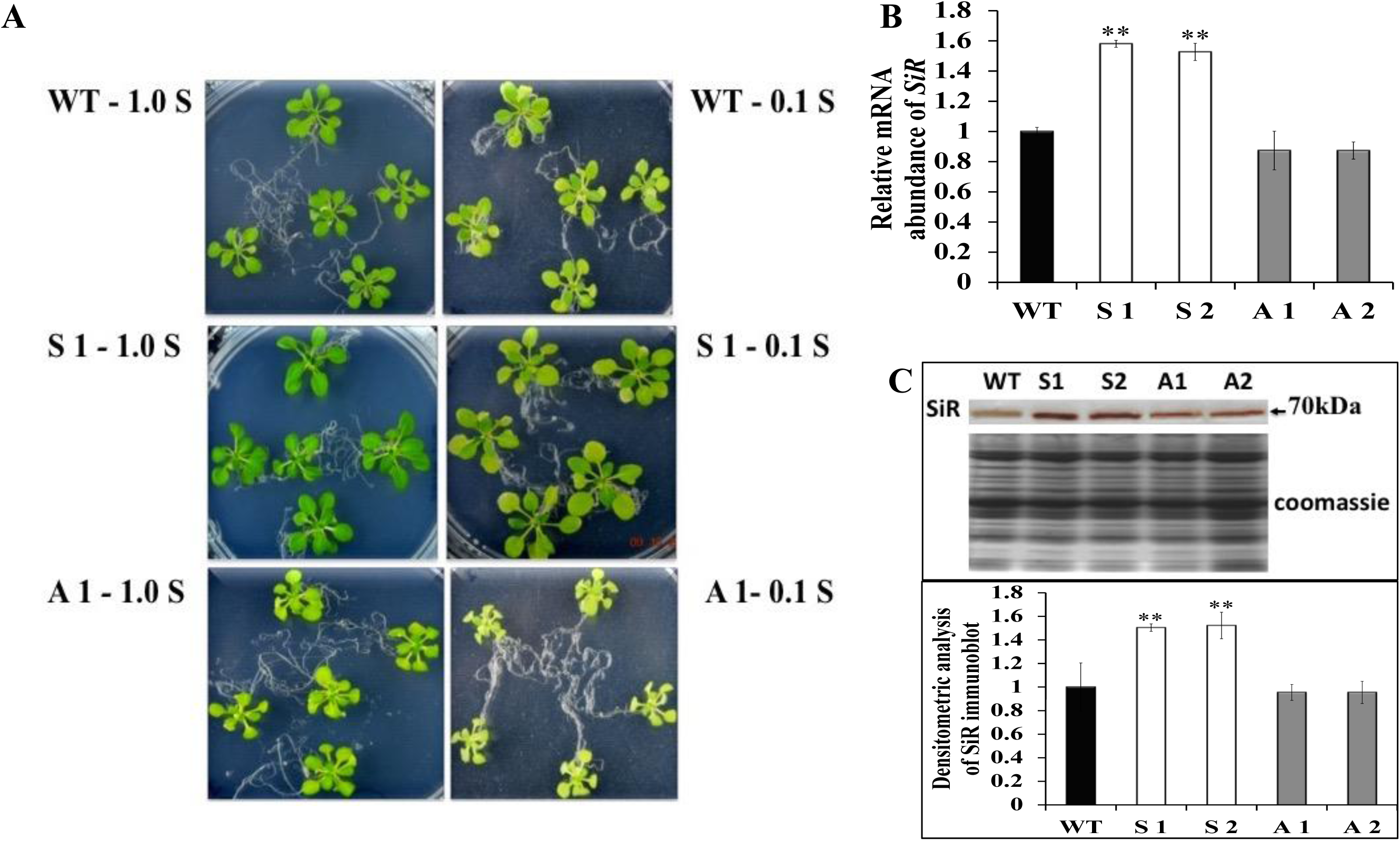
Responses of *WT and AtSirB* transgenic plants to sulfur limitation. WT and *AtSirB* (S 1 and A 1) plants were grown at 22^0^C under 14h L / 10h D photoperiod in a light intensity of 100 µmoles photons m^−2^ s^−1^ for 15 days in optimal sulphur media (1S) and subsequently transferred into minimal sulfur (0.1S) media for five days. Concentration of Sulphur in normal media (1S) was 2mM. (A) Phenotype of WT and *AtSirB* transgenic plants in S-limitation condition, (B) the *SiR* gene expression measured by qRT-PCR and (C) protein abundance measured by Western blot using antibodies against sulfite reductase (*Zea maize*). Standard deviation is shown as the error bar. WT, S1, S2, A1, and A2 significantly differ from each other as indicated by the (*) asterisk (P< 0.05).

**Figure 8:**
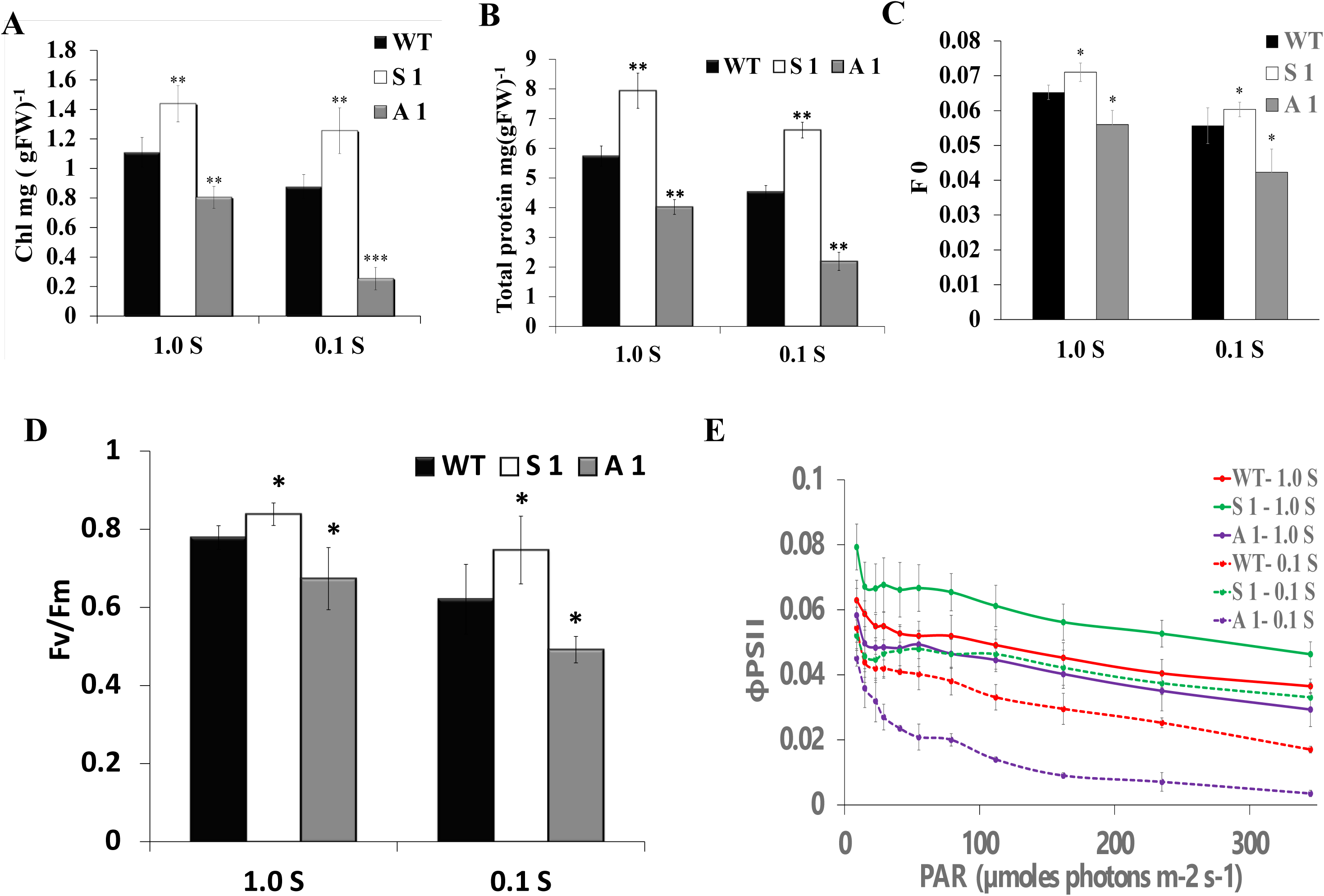
Changes in physiological parameters in S-limitation condition. WT and *AtSirB* transgenic plants (S1 and A 1) were grown at 22^0^C under 14h L / 10h D photoperiod in a light intensity of 100 µmoles photons m^−2^ s^−1^ in MS for 15 days and subsequently transferred for 7 days into Sulfur limitation condition (0.1S) and their Chl, protein and Chl fluorescence parameters were monitored. (A) Total Chl, (B) Total Protein, (C) F0, (D) Fv/Fm, (E) фPSII. An average of 10 replicates were used for Chl and protein content and 10 replicates were used for Chl a fluorescence parameter. The standard deviation is shown as error bar. WT, S1, and A1 significantly differ from each other as indicated by the (*) asterisk (P< 0.05).

### Photosynthetic response of WT and transgenic plants grown in S-limitation condition

WT and transgenic plants were raised for 15 days in normal MS before being transplanted to agar plates having Hoagland medium, which had its S content adjusted to either an ideal (1.0S) or deficient (0.1S) level by lowering SO_4_^−2^ by substituting with Cl-. The Fo, Fv/Fm, and фPSII were measured after seven days of development in the S-limitation medium. Fo declined in S-limiting condition in WT, S1, and A1 plants, although S1 plants had higher F0 than WT, whereas A1 had lower F0 than WT at all nitrogen levels **(Figure 8C)**. The фPSII rose in response to PAR. However, the rise in фPSII of S1 plants was higher than WT in limiting and saturating light intensities. Similarly, in comparison to WT, S1 plants had a greater Fv/Fm ratio, whereas in antisense plants, the Fv/Fm ratio decreased **(Figure 8D)**. In S starvation media, an increase in the фPSII of S1 plants over WT was highly pronounced in limiting and saturating light intensities.

The increased slope of фPSII at limiting light intensities suggests better energy capture by the light-harvesting complex, and the higher light-saturated rate implies an increase in the amounts of electron transport components in S1 plants. In A1 plants, the фPSII decreased in high light intensities in S-deficient media **(Figure 8E)**.

## Discussion

Although C assimilation is a well-known light-dependent reaction, N and S assimilation is indirectly regulated by light. The reducing side of PSI serves as a control point for the mutual regulation of the NO_2_^−^ and CO_2_^−^ assimilation processes, which interact at the level of photosynthetic electron transport. Ferredoxin-NADP reductase and nitrite reductase effectively compete for decreased ferredoxin. NO_2_^−^ reduction facilitates the synthesis of ATP for alternate purposes, including the synthesis of proteins and carbohydrates in chloroplasts. The gene expression of nitrate reductase is regulated by light via phytochromes (Sharma & Sopory, 1984; Sherameti et al., 2002). The *AtSirB is* responsible for N and S assimilation in plants, and these processes are dependent on light for proper utilization of N and S compounds taken up from the soil (LaBrie and Crawford, 1994; Bork et al., 1998). We have shown that the *AtSirB* is a light-regulated gene that is present in a minimal amount in etiolated tissues and highly upregulated after light exposure due to light-responsive elements in its promoter (Garai et al., 2016). It may be inferred that similar to the light-regulated expression of apoenzyme, the gene expression of the enzyme required to synthesize the cofactor is also photo-regulated (Becker, Foyer, and Caboche, 1992; Garai et al., 2016; Rajasekhar and Mohr, 1986).

Our study on overexpression of AtSirB working in the branched pathway of plant tetrapyrrole biosynthesis responsible for the biosynthesis of siroheme that governs N an S assimilation demonstrates the co-modulation of NiR and SiR responsible for amino acid and protein synthesis. We have partly diverted uroporphyrinogen III, an intermediate involved in the biosynthesis of Chl, a central pigment of carbon assimilation, for increased N and S assimilation. qRT-PCR results show that *AtSirB* expression increased in *AtSirBx* plants and declined in antisense plants. Similarly, protein expression increased in *AtSirBx* lines and reduced in antisense lines. In comparison to WT*, AtSirBx* plants were larger and greener in color. Most of the *AtSirB* antisense plants were quite pale and could not survive; this could be due to severe silencing of *SirB* resulting in highly impaired NiR and SiR functions and consequent accumulation of toxic metabolites, i.e., nitrite and sulfite (Bork et al., 1998; Takahashi et al., 2001). The severe impairment of N and S assimilation also contributed to plant death. Only a few antisense lines survived, though they looked pale and were smaller than WT. Compared to WT, the number of lateral roots per plant was higher in *AtSirBx* plants. However, the antisense plants had the same number of lateral roots as the WT plants. In contrast to antisense plants, the fresh and dry weights of *AtSirBx* were higher than WT’s. It could be because of overexpressers assimilating N, S, and C at higher rates. The present study reveals that the reduction of NiR activity in *AtSirB* antisense plants down-regulates NR reaction to avoid excess accumulation of toxic nitrite. The downregulated NR function could reduce nitrate uptake from the growth medium by suppressing the root-localized high-affinity nitrate transport system (HATS). Therefore, plants would stifle the development of nitrate-absorbing lateral roots. It is further supported by Little et al. (2005), who showed that a defect in HATS stimulates lateral root initiation in *Arabidopsis*.

Conversely, the total protein of antisense plants declined by 18%. This resulted from a drop in NiR activity brought on by a decrease in siroheme biosynthesis. The NR activity also declined to prevent the accumulation of toxic NO_2_^−^. Although NiR apoprotein content was not significantly affected in antisense lines, due to reduced availability of the cofactor siroheme, the NiR activity declined by 60%. The siroheme is not directly required for NR activity; however, the NO_3_^−^ reduction dropped by 23%. It demonstrates the co-regulation of NR and NiR; in both over-expresser and antisense plants, there is still a strong probability that *NR* and *NiR* gene expression and activity are co-regulated. The *NR* and *NiR* transcript levels are known to be co-induced by NO_3_^−^ and light (Faure et al., 1991). However, it is possible that siroheme post transcriptionally modulates NR.

In the present experimental conditions, NO_3_^−^ and the light were optimally present, which rules out the possibility of inherent down-regulation of their gene expression in the antisense plants. Although NiR activity declined by 60% in antisense plants, the total protein content decreased by 18%. It suggests that plants may partially overcome the diminished NiR reaction, which creates the NH_4_^+^ essential for synthesizing amino acids and proteins in an optimal NO_3_^−^ containing growing medium. Conversely, *AtSirBx* plants having higher NiR activity had enhanced NR function to supply the substrate NO_2_^−^ for the augmented NiR reaction in the overexpressor stimulates NR and, therefore, could encourage nitrate uptake by promoting lateral root growth (Little et al., 2005).

Chl synthesis is one of the most significant cellular processes for plants’ photosynthesis. Chl content increased significantly in *AtSirBx* lines and decreased in antisense lines. In *AtSirBx* lines, NiR gene (*Nii*) and protein expression increased, which resulted in increased NiR activity. The increase in NiR activity was due to the enhanced availability of the apoprotein and the prosthetic group siroheme in *AtSirBx* plants. In consonance with augmented NiR and NR activities, the protein content of the over-expressers was higher than that of the WT. Our results demonstrate the co-regulation of NiR and NR gene expression and their activities that predominantly contribute to N assimilation in non-leguminous plants. In consonance with the increase in the protein content, the Chl content of *AtSirBx* plants was also higher.

Earlier attempts to increase the protein content of plants by genetically modulating nitrate transporters, NR, NiR, GS, and GOGAT, did not succeed. Plants overexpressing the high-affinity nitrate transporter (HATS) gene *NpNRT2;1*, despite having an increased NO3-influx to the roots, did not have higher NUE (Fraisier et al., 2000). These results imply that increased nitrate uptake via genetic engineering may not necessarily improve NUE. *Nia1*, which codes for NR, was overexpressed constitutively in tobacco to improve NUE, which resulted in a drop in nitrate levels in tobacco and potatoes (Quilleré et al., 1994, Djennane et al., 2002). No discernible difference in biomass or protein accumulation was seen in NR overexpressers despite having more NR protein available (Vincentz and Caboche, 1991). Over-expression of *NiR* with 35S promoter in *N. plumbaginifolia* and *Arabidopsis thaliana* results in higher NiR activity of transgenic plant grown in nitrate-supplemented media; however, there was no increase in total plant protein nor any change in plant phenotype (Crété et al., 1997; Takahashi et al., 2001).

Therefore, it is yet uncertain if the over-expression of NR/NiR will significantly increase NUE. However, it is possible that different crops could respond differently. Similarly, attempts to improve NUE by changing the plastidic GS2 and Fd-GOGAT genes have had only patchy results. Over-expression of GS2 in transgenic tobacco plants has increased high-light tolerance and improved photorespiration (Kozaki and Takeba, 1996). Over-expression of plastidic GS2 with leaf-specific *RSS* promoter in tobacco resulted in improved assimilation of ammonia into amino acids (Ferrario-Méry et al., 2002). However, it was not accompanied by a rise in the total protein contents of plants. Overexpression of cytosolic glutamine synthetase (*GS1*) in different crops grown in various nitrate concentrations indicated a limited increase in NUE (Man et al., 2005; Fei et al., 2006). In our present study, the NiR’s assembly and activity may be significantly influenced by the enhanced siroheme prosthetic group synthesis in *AtSirBx* plants.

In *AtSirB* over-expressers, improved availability of siroheme increased NiR activity and improved S assimilation by NiR and SiR, respectively. Siroheme is limited, and its increased synthesis will likely enhance the NUE. Therefore, increased synthesis of ammonium and sulfide could have stimulated amino acids and protein synthesis. Increased protein content and upregulated photosynthesis resulted in a bigger phenotype and higher biomass accumulation in *AtSirBx* plants. It demonstrates that the NUE is improved when the cofactor synthesis is stimulated rather than the buildup of apo-proteins of NiR or SiR **(Figure 9).** Our previous study with the other siroheme biosynthesis gene *AtUPM1* has demonstrated that when it is overexpressed in *A. thaliana*, it increases amino acids and protein content in transgenics (Garai and Tripathy, 2018).

**Figure 9:**
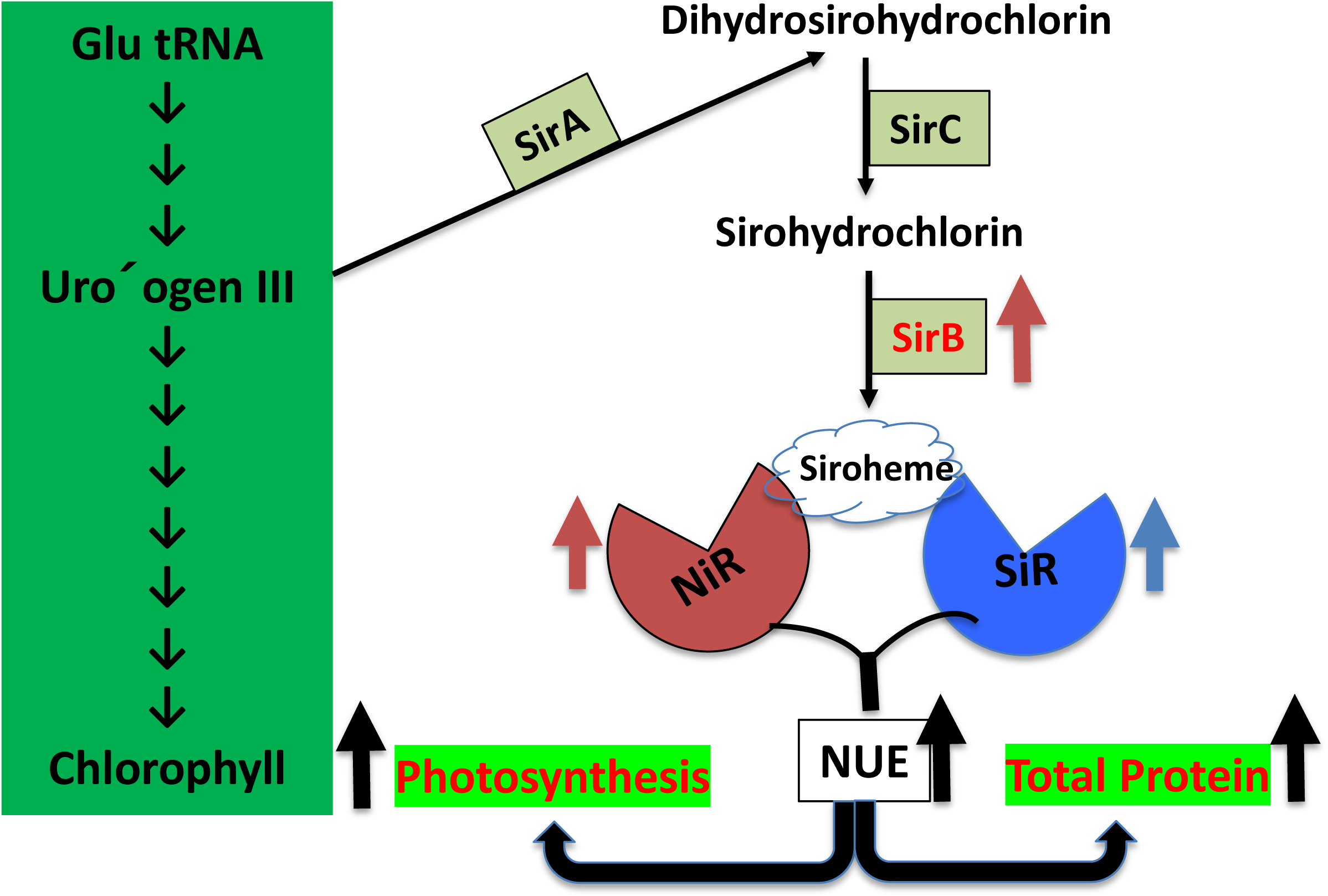
A cartoon of *SirB* expression in Arabidopsis. Overexpression of SirB (Sirohydrochlorin ferrochelatase) increases the abundance of Siroheme in over-expresser plants that further induces expression/activity of nitrite reductase and sulfite reductase in over-expressers. Induction of NiR and SiR enhances the nutrient use efficiency (NUE) of over-expresser plants. Enhancement of NUE increases total protein content which ultimately increases the photosynthetic efficiency (PE) of *AtSirB* overexpressing Arabidopsis plants.

### Overexpression of *AtSirB* protects plants from N deficiency

In order to test whether WT and *AtSirB* transgenic plants could tolerate N deficiency, these plants were grown in N-limiting media. Due to decreased Chl accumulation, the WT plants nearly blanched due to N constraint (0.1N). The *AtSirBx* plants were greener than the WT and had more Chl than WT under similar growth conditions. However, in *AtSirB* antisense plants, the N limitation caused severe phenotypical changes, including bleaching at 0.1N due to severe reduction of protein and Chl contents. Synechocystis and Chlorella grown in N-deficient media had decreased photosynthetic activity, smaller antennas, and slower electron transport rates, reduced C-dependent oxygen evolution and biomass (Salomon et al., 2013; Kumari et al., 2021) that supports our present data. The total protein content was higher in *AtSirB*x plants than WT plants and reduced in *AtSirB* antisense plants. Under normal (1N) N level, the protein content of over-expresser plants was 35% higher, and in antisense, it was lower by 11% than WT. *AtSirBx* plants had a substantially higher protein content (95%) than WT plants under N-limiting circumstances. Conversely, the antisense plants had a lot lower (32%) proteins due to N starvation than the optimal growth medium. The decreased protein content of plants grown in nitrate-deficient media was due to reduced N assimilation caused by lower NiR and NR activities. Similar to their activities, *NR* and *NiR* gene expression increased in *AtSirBx* plants in optimal growth media. The photosynthetic efficiency of the *AtSirBx* plants was better than those of WT in optimal growth medium (1N) due to the higher protein levels. On the other hand, the photosynthetic performance of *AtSirB* antisense plants decreased due to their reduced protein contents.

The values of фPSII in light-adapted plants are higher in transgenics than WT both in limiting and saturating light intensities, suggesting better light utilization likely due to an increase in the abundance of light-harvesting complex and better light utilization of absorbed energy because of robust PSII. Conversely, the фPSII in light-adapted plants were lower in antisense plants than the WT both in low and high light intensities, suggesting reduced light absorption and utilization likely due to lower abundance of light-harvesting complex and reduced light utilization of absorbed energy by frail PSII.

The Fv/Fm ratio decreased in WT, sense, and antisense plants on N-starvation media. However, the *AtSirBx* plants consistently showed higher Fv/Fm ratios than WT at all N levels, while the *AtSirB* antisense plants always had smaller Fv/Fm ratios. The increased protein content of *AtSirBx* plants contributed to the better development of the thylakoid membranes even under N-limiting conditions. In normal and N-deficient conditions, the фPSII was higher in *AtSirBx* and lower in antisense plants in all measured light intensities. In normal growth media, the Fv/Fm value is essentially constant but declines in medium with nutrient constraints, which can be used to identify nutritional limitations (White et al., 2011; Zhang et al., 2013). In the WT and *AtSirBx* plants, the фPSII increased in response to a rise in PAR in N-sufficient and N-deficient media, albeit to a larger extent in the over-expressers. In *AtSirBx* plants, the greater light-saturated rate denotes an increase in the components of the electron transport chain. In contrast, the increased slope of фPSII at limiting light intensities shows improved energy collection by the larger light-harvesting complex. The фPSII decreased in antisense plants in low and high light intensities in all N levels, indicating a decline in energy capture and usage due to decreased light-harvesting antennae and reduced abundance of the electron transport components.

### Overexpression of *AtSirB* protects plants from S deficiency

Numerous aspects of S metabolism have been well explored. Sulfate is absorbed and is reduced to sulfite. The sulfite is further reduced to sulfide by 6 electron reduction mediated by chloroplastic sulfite reductase (Leustek and Saito, 1999). *Arabidopsis* has five functional subgroups of sulfate transporters (Smith et al., 1995). Sulfate deficiency increases the development of these high-affinity sulfate transporters, which help the soil absorb sulfate when sulfate is scarce (Maruyama-Nakashita et al., 2003; Takahashi et al., 1997). These sulfate transporters activate the sulfate uptake from the soil in S-deficiency conditions, but when plants cannot absorb enough sulfate, the decreased sulfate uptake results in decreased S assimilation activity (Yoshimoto et al., 2007). Plant tissues eventually contain less S due to the limited amounts of S available in soil and plants (Kutz et al., 2002; Prosser et al., 2001). Inhibition of sulfate assimilation results in decreased cysteine and methionine, total protein, and Chl content of plants (Nikiforova et al., 2003). Similarly, our results demonstrated that in an S-deficient medium (0.1S), the WT plants had very low Chl and were pale green after seven days of growth. However, the *AtSirBx* plants, because of better utilization of available low amounts of S, had higher Chl and were greener than WT. Under S-starvation, the *AtSirB* antisense plants had severe phenotypical changes; plants became severely blanched due to reduced Chl contents. S-starvation causes a rise in reactive oxygen species (ROS) (Shin et al., 2005).

*AtSirBx* plants had increased SiR protein abundance over that of WT. In agreement with protein expression*, SiRB* mRNA expression substantially increased in *AtSirBx* plants compared to WT. In antisense plants, *SiR* mRNA abundance was slightly reduced. Antisense plants had no significant change in abundance of SiR apoprotein as an adaptive response to the minimal availability of siroheme for the holoenzyme assembly. In the plastid ferredoxin-dependent, SiR converts sulfite to sulfide to synthesize cysteine and methionine. *AtSirBx* plants could produce more proteins than WT because of the increased availability of S-containing and non-S-containing amino acids. However, protein content was lower in antisense plants, likely due to decreased NiR and SiR activity. *AtSirBx* plants had better photosynthetic rates than WT plants in optimal growth medium (1S) due to the higher protein content. Conversely, the photosynthetic capacity of antisense plants is reduced because of their lower protein concentrations. The Fv/Fm, a measure of the plant’s quantum efficiency in the dark-adapted plants, declined when WT, *AtSirBx,* and antisense plants were grown on an S-starvation medium. The antisense plants had a lower Fv/Fm than the WT, whereas the *AtSirBx* plants consistently had a higher Fv/Fm ratio under both S-optimal and S-deficient growth conditions. This was due to increased protein contents of *AtSirBx* plants that contributed to the better development of the thylakoid membranes in S-starvation growth conditions. The фPSII in *AtSirBx* plants was higher under normal and S-deficient conditions than in antisense plants. The фPSII rose in response to PAR; the *AtSirBx* plants had higher фPSII in both low and high light intensities, demonstrating that the light-harvesting complex and the electron transport components of PSII were higher in the over-expressers. In S-deficiency, the фPSII declined in WT and, to a larger extent, in antisense plants, suggesting loss of their PSII antennae and electron transport components. In S-deficient AtSirBx plants, the фPSII was much higher than in S-starved WT plants, suggesting the preservation of photosynthetic apparatus and tolerance to S-deficiency.

## Conclusion

Excessive use of fertilizers is having a devastating impact on the environment. Increased N use efficiency of the crop will decrease fertilizer burden on the soil for increasing productivity. Our data suggest the possibility of increasing N-use efficiency and photosynthesis and overcoming soil N– and S-deficiency by the overexpression of *SirB* in crop plants for sustainable agriculture.

## Materials and Methods

### Plant materials and growth conditions

The *Arabidopsis thaliana* Columbia-0 (Wild-type: WT) and *AtSirB*-overexpressing lines (*AtSirBx*, S1& S2) and silencing lines antisense (A1&A2) were cultivated in a growth chamber (Conviron, Canada) (100 μmoles photons m^−2^s^−1^), and a 14 h light/ 10 h dark photoperiod at 22^0^C (Pattanayak et al., 2005).

### Cloning and transformation of AtSirB in Arabidopsis

*AtSirB* was amplified from *Arabidopsis* cDNA from mature leaf, with the help of gene-specific forward (FP) and reverse (RP) primers were made according to *AtSirB* cDNA sequence **(Supplementary Table. 1)**. For the amplification, a PCR program was designed having 94°C for 5 minutes followed by 30 cycles of 94° C (denaturation) for 30 seconds, 56° C (annealing) for 30 seconds, 72^0^ C (extension) for 45 seconds using a standard PCR reaction set up. The resulting amplification product was gel purified and ligated to a pGEMT-easy (Promega, USA) vector. The recombinant plasmid (pGEMT-easy-*AtSirB*) was transformed into competent *E*.*coli* (DH5α) cells. Plasmid DNA was prepared, and the nucleotide sequence of the *AtSirB* was confirmed by sequencing using a standard procedure (Sambrook et al., 1989). The modified pCAMBIA1304 binary vector was used to clone the full-length AtSirB cDNA from the recombinant pGEMT-Easy plasmid after it had been EcoRI digested and the product had been oriented in both sense and antisense. *Arabidopsis thaliana* (Col-0) was transformed using the vacuum infiltration method after the pCAMBIA1304: *AtSirB* construct was transformed to the *Agrobacterium tumefaciens* strain GV3101 (Clough and Bent, 1998; Das, 2011). Primary transformants were cultivated until T3 generation after being screened in half-MS containing 50 mg/L kanamycin to establish homozygous lines for further study. The sense orientation of *AtSirBx* plants was done by gDNA PCR using 35S Int-FP and *AtSirB* RP, whereas the antisense orientation of antisense transgenic plants was tested using 35S Int-FP and *AtSirB* FP.

### Quantitative RT-PCR analysis to check AtSirB mRNA expression in AtSirB transgenic plants

TRIzol reagent (Invitrogen, USA) was used to isolate total RNA from 15-day-old *Arabidopsis* seedlings. To ensure no DNA was present, the RNA was processed with RNase-free DNase I (Promega, USA). In a 50 μL reaction volume, 2 μg of total RNA, an oligo (dT) primer, and AMV reverse transcriptase (Promega, USA) were used to produce cDNA for reverse transcription (RT) PCR (Pattanayak and Tripathy, 2011). The cDNA was diluted 1:10 and utilized as a template for qRT-PCR reaction with 2μL of the diluted cDNA. The gene-specific primers mentioned in **Supplementary Table 1** were used in PCR reactions per the manufacturer’s instructions using an Applied Biosystems (Life Technologies, Carlsbad, CA) real-time PCR system. Using the comparative CT approach (Schmittgen and Livak, 2008), expression was normalized to Actin. The resulting data are shown as the standard deviation and relative means of three biological replicates.

### Antisense mRNA Expression in antisense AtSirB plants

Gene-specific primers were employed to prime first-strand cDNA synthesis rather than Oligo (dT) or random hexamers. Reverse transcription from RNA obtained from *AtSirB* transgenic strains was done using the forward primer to look for antisense mRNA. Only the antisense mRNA is bound by the forward primer. As a result, if there is no antisense mRNA, no cDNA synthesis will occur from the forward primer (Song and Wang, 2009). The result demonstrates that the *AtSirB* gene product was amplified when PCR was performed using the cDNA prepared from gene-specific forward primer only. cDNA prepared using total RNA from wild-type plants was used as a template for another set of PCR that served as negative control.

### Total protein quantification from AtSirB transgenic seedlings

To calculate the total protein content, seedlings were homogenized with liquid N. The homogenized tissues were mixed with 1 ml of an isolation buffer of 56 mM Na_2_CO_3_, 5 mM DTT, 10 mM isoascorbate, 12 % sucrose, two mM EDTA, and % SDS. The samples were adequately vortexed after being transferred from the homogenate to the Eppendorf tube. Samples were heated at 80^0^C for 20 minutes, followed by centrifugation at 13000 rpm for 5 minutes. A new Eppendorf tube was used to collect the supernatant. The protein content in various extracts was estimated using Bradford’s method (Bradford, 1976). A polyclonal antibody produced against the AtSirB protein was used to blot approximately 10 μg of the total protein supernatant separated onto SDS PAGE (Laemmli, 1970). Monoclonal antibody against NiR (nitrite reductase) was prepared commercially, and SiR (Sulfite reductase) antibody (*Zea mays*) was kindly donated by Toshiharu Hase, Institute for Protein Research, Osaka University, Japan.

### Isolation of thylakoid membranes

Isolation of thylakoid membrane was done by the method described in ***Supplementary methods*.**

### Chl estimation in AtSirB transgenic

Chl was extracted from tissues and thylakoid membranes in a safe green light. Leaf samples (as well as seedlings) were homogenized in 10 ml of 90% cold ammonical acetone using a previously chilled mortar and pestle. For each batch, more than ten replicates were collected. Homogenate was centrifuged at 4°C for 10 minutes at 10,000 rpm. The absorbance of the supernatant was measured at 663 nm, and 645 nm to calculate total Chl. Chl was estimated according to (Porra et al., 1989).

### Extraction and determination of tetrapyrroles

Extraction and determination of tetrapyrroles was done by the method described in ***Supplementary Methods*.**

### Pulse amplitude modulation (PAM) measurements in AtSirB transgenics

All quantifications of Chl fluorescence were made using a PAM-2100 fluorometer (Walz, Effelteich, Germany) as described in ***Supplementary Methods***.

### Measurement of morphological parameters in AtSirB transgenics

WT and *AtSirB* transgenic plants were raised for three weeks in MS medium. The fresh weight of the plants was measured after they were removed from Petri plates and placed on tissue paper to absorb moisture. Whole plants were wrapped in aluminium foil and baked at 70^0^C for 48 hours to determine their dry weight. Primary root length was measured with Image J software, and the numbers of lateral roots were counted individually under a light microscope.

### Quantification of Nitrite reductase and Nitrate reductase activity in AtSirB transgenic Nitrite reductase assay

Nitrite reductase activity assessed as the decline of nitrite ion in the assay mixture as described by (Takahashi et al., (2001). For details protocol refer to ***Supplementary Methods*.**

### Nitrate reductase assay

NR activity was measured as described by Kaiser and Lewis (1984), with a minor change to determine NO_2_^−^ ion formation in the reaction volume. For details refer to ***Supplementary Methods***.

### AtSirB transgenic plants in N –starvation medium

Plants were grown photo-periodically (16h L/8h D) at 22^0^C in 100 μmoles photon m^−2^ s^−1^ light intensity for 15 days in normal MS medium. After 15 days, plants of equal size were taken and transferred to optimal (1N) or N deficient (0.1N) Hoagland media in an agar plate (**Supplementary Table. 2)**. After seven days of growth in N-limiting medium, WT and *AtSirB* sense and antisense plants were analyzed. Chl, Chl *a* fluorescence (PAM), protein, and NiR activities were measured.

### Accumulation of Starch in AtSirB transgenic plants in N starvation

Plants were grown photo-periodically (16h L/8h D) at 22^0^C in 100 μmoles photon m^−2^ s^−1^ light intensity for 15 days in normal MS medium. After 15 days, plants of equal height were transferred to optimal or N-deficient media prepared in an agar plate with a Hoagland solution as in **Supplementary Table. 2**. After ten days of growth in N-starvation medium, WT, AtSirB sense, and antisense plants were kept in methanol for 24 hours. Subsequently, plants were stained with 0.2% w/v of iodine solution that developed a blue color due to starch accumulation in the leaves in N-deficient plants (Hermans et al., 2016).

### AtSirB transgenic plants in S –starvation medium

WT and *AtSirB* transgenic plants were grown photo-periodically (16h L/8h D) at 22^0^C in 100 μmoles photon m^−2^ s^−1^ light intensity for 15 days in normal MS medium. After 15 days, plants of equal size were taken and transferred to optimal (1S) or S deficient (0.1S) Hoagland media in an agar plate **(Supplementary Table. 2)**. After seven days of growth in S-starvation medium, WT, and *AtSirBx* and antisense plants were analyzed. Chl, carotenoids, Chl *a* fluorescence (PAM), and protein contents were measured.

## Acknowledgment

The authors thank the Department of Biotechnology, Govt. of India for financial assistance.

## Author statement

NCJ performed the experiments, and BCT conceptualized the research work in the manuscript.

## Figure Legend

**Supplementary Figure S1:** Root Morphology *AtSirB* transgenic. WT and *AtSirb* Sense (S1) and antisense (A1) plants were grown vertically at 22^0^C under 14h L / 10h D photoperiod in a light intensity of 100 µmoles photons m^−2^ s^−1^ for 3 weeks in MS medium.

**Supplementary Figure S2:** Protochlorophyllide, protoporphyrin IX content in WT and *AtSirB* transgenic plants-. WT and *AtSirB* (S1 and A1) plants were grown at 21^0^C under 14h L / 10h D photoperiod in a light intensity of 100 µmol photons m^−2^ s^−1^ for three weeks in MS medium and their (A) Chl biosynthetic intermediate protochlorophyllide and (B) protoporphyrin IX were measured as described in “Materials and Methods”. An average of 10 replicates are used for each data point for protochlorophyllide and protoporphyrin IX measurements. Standard deviation is shown as the error bar. WT, S1, and A1 significantly differ from each other as indicated by the (*) asterisk (P< 0.05).

**Supplementary Figure S3:** Sucrose accumulation as signature of N-starvation. WT and *AtSirBx* and antisense plants were raised in MS medium for 15 days and subsequently sifted to different concentration of Nitrate (0.1N of normal MS concentration) for 7 days were taken and iodine staining was done to check the translocation of starch.

